# Fungal jasmonate as a novel morphogenetic signal for pathogenesis

**DOI:** 10.1101/2021.04.04.438374

**Authors:** Yingyao Liu, Martin Pagac, Fan Yang, Rajesh N. Patkar, Naweed I. Naqvi

## Abstract

A key question that has remained unanswered is how pathogenic fungi switch from vegetative growth to infection-related morphogenesis during a disease cycle. Here, we identify a fungal oxylipin analogous to the well-known phytohormone jasmonic acid, as the principal morphogenesis signal responsible for such a developmental switch to pathogenicity in the rice-blast fungus *Magnaporthe oryzae*. We explored the molecular function(s) of such intrinsic jasmonic acid during pathogenic differentiation in *M. oryzae* via *OPR1*, which encodes a 12-Oxo-phytodienoic Acid Reductase essential for its biosynthesis. Loss of *OPR1* led to prolonged vegetative growth, and a delayed initiation and improper development of infection structures in *M. oryzae*, reminiscent of phenotypes observed in mutants (e.g. *pth11*Δ and *cpka*Δ) that are compromised for cyclic AMP signaling. Genetic- or chemical-complementation completely restored proper germ tube growth and appressorium formation in *opr1*Δ. Liquid chromatography mass spectrometry-based quantification revealed increased OPDA accumulation and a significant decrease in JA levels in the *opr1*Δ. Most interestingly, exogenous jasmonic acid also restored appressorium formation in the *pth11*Δ mutant that lacks G protein/cyclic AMP signaling. Epistasis analysis placed fungal jasmonate upstream of the cyclic AMP signaling in rice blast. Lastly, we show that intrinsic jasmonate orchestrates the cessation of vegetative phase and initiates pathogenic development via a regulatory interaction with the cyclic AMP cascade and redox signaling in rice blast.

## Introduction

Phytohormone mimics produced by some pathogenic fungi participate in the cross-kingdom communication between fungi and hosts for their benefits. This striking phenotype has become a rapidly emerging research field (Kazan and Lyons 2014, Patkar and Naqvi 2017). We found that jasmonic acid (JA) and its derivate 12OH-JA are synthesized by the rice blast fungus *Magnaporthe oryzae* and subsequently secreted into the host cells to suppress the JA-dependent plant immunity (Patkar, Benke et al. 2015). This remarkable discovery also identified that the plant pathogenic fungus, *Magnaporthe oryzae*, follows *Lasiodiplodia theobromae* (Miersch, Preiss et al. 1987), *Fusarium oxysporum* and *Gibberella fujikuroi* (Miersch, Günther et al. 1993), in biosynthesis of such fungal JA-like oxygenated lipids or oxylipins *in vivo*.

JA is a crucial plant defense hormone and ubiquitously distributed in land plants. The biosynthesis pathway of jasmonate and its physiological functions in plants have been well studied. Briefly, polyunsaturated fatty acids, such as α-LeA, are synthesized from a membrane component and secreted into plastids (Ishiguro, Kawai-Oda et al. 2001). Catalyzed by 13-Lipoxygenases (13-Lox), Allene oxide synthase (Aos), and Allene oxide cyclase (Aoc) subsequently (Feussner and Wasternack 2002, Park, Halitschke et al. 2002, Schaller, Zerbe et al. 2008, Shin, Van et al. 2008, Hughes, De Domenico et al. 2009, Dave and Graham 2012, Christensen, Huffaker et al. 2015), the key intermediate product *cis*-OPDA is generated in the chloroplasts (Stintzi 2000, Park, Halitschke et al. 2002, Hughes, De Domenico et al. 2009, Dave and Graham 2012, Kombrink 2012). Upon the migration of *cis*-OPDA to peroxisomes, it is reduced by Opr3, and then its carboxylic acid side chain is shortened via three cycles of β-oxidation reactions (Li, Schilmiller et al. 2005, Schilmiller, Koo et al. 2007). Finally, *cis*-jasmonic acid is biosynthesized in the peroxisomes, and trafficked to the cytosol where it can be modified into different kinds of jasmonates (Wasternack and Strnad 2018). Although the mechanism remains unclear, recent studies suggest that jasmonate and/or its derivates are also secreted out of the cells (Patkar et al. 2015).

Compared to the knowledge in plants, the synthesis and metabolism of jasmonate-like oxylipins in fungi remain largely unexplored. For instance, OPDA, the critical intermediate in JA biosynthesis, has been identified in *L. theobromae* and *F. oxysporum*, the relevant metabolic enzymes along with the synthesis pathway, such as Allene oxide synthase (Aos) and OPDA reductase (Opr), have not been characterized in detail (Tsukada, Takahashi et al. 2010, Chen, Jernerén et al. 2017, Matsui, Amano et al. 2017).

One of the most challenging issues in *in silico* investigation of microbial jasmonate biosynthesis has been the lack of proper sequence conservation in orthologous proteins in fungi compared to the plant counterparts. For example, as a member belonging to the cytochrome P450 (CYP) family, the fungal Aos in *Aspergillus terreus* has 38% amino acid identity with the plant 5,8-LDS but shares only 25% with CYP8A1 or CYP74A (Hoffmann, Jernerén et al. 2013). The Aos in *F. oxysporum* is a 9-Dox-Aos fusion protein with an N-terminal dioxygenase (Dox) domain and and an Aos motif at the C terminus. Although the sequence alignment shows that this C terminal AOS is homologous to the AOS of *A. terreus* with over 50% identity, the 9-Dox-Aos in *F. oxysporum* is not the biosynthesis-related enzyme of jasmonic acid, as the final products are α-Ketos and γ-Ketos (Hoffmann and Oliw 2013).

Another central question that has remained unexplored is: What are the physiological functions of jasmonate(s) in fungi? Thus far, the investigation of fungal jasmonate function has predominantly focused on the plant immune response. For example, the most well-elucidated microbiota-jasmonate mechanism is Coronatine (COR) in the pathogenic bacterium *Pseudomonas syringae* (Misaghi 1982). COR serves as the analogue of the bio-active JA-Isoleucine and competes with binding to the co-repressor Coi1, which results in blocking of JA-based plant defence (Alarcón-Chaidez, Penaloza-Vázquez et al. 1999). Lasiojasmonate (LasA), an inactive jasmonate derivative produced by *L. mediterranea*, accumulated *in vivo* served as a JA pool to be catalysed into JA-Ile. Resultant JA-Ile secreted into host cells at later infection stage to activate plant JA-related signalling via Coi1-JAZ dependent pathway and cause host cell death consequently (Chini, Cimmino et al. 2018). Nevertheless, the physiological role of intrinsic JA in these fungal systems remains unclear.

The Blast pathosystem, comprising Rice and *Magnaporthe oryzae*, represents one of the most destructive fungal diseases of cultivated cereal crops. *M. oryzae* is a hemibiotroph that initially keeps the host cells alive prior to switching on the destructive necrotrophic cell death modality therein (Patkar et al 2015). Conidia/asexual spores of the blast fungus follow an intricate developmental program in response to specific surface cues to differentiate into specialized infection structures, the appressoria, at the tips of emerging germ tubes (**Fig. 1A**; Kou and Naqvi 2016). In the absence of such cues/inductive surfaces, the germ tubes continue vegetative development in a unipolar fashion and produce mycelia and subsequently the conidia. An enormous turgor pressure generated within the appressorium is utilized to physically rupture the host cuticle, and a thin penetration peg enables entry into the host (Kou and Naqvi 2016). The peg further differentiates into bulbous infectious hyphae that colonize host tissues resulting in typical blast symptoms.

**Figure 1.**
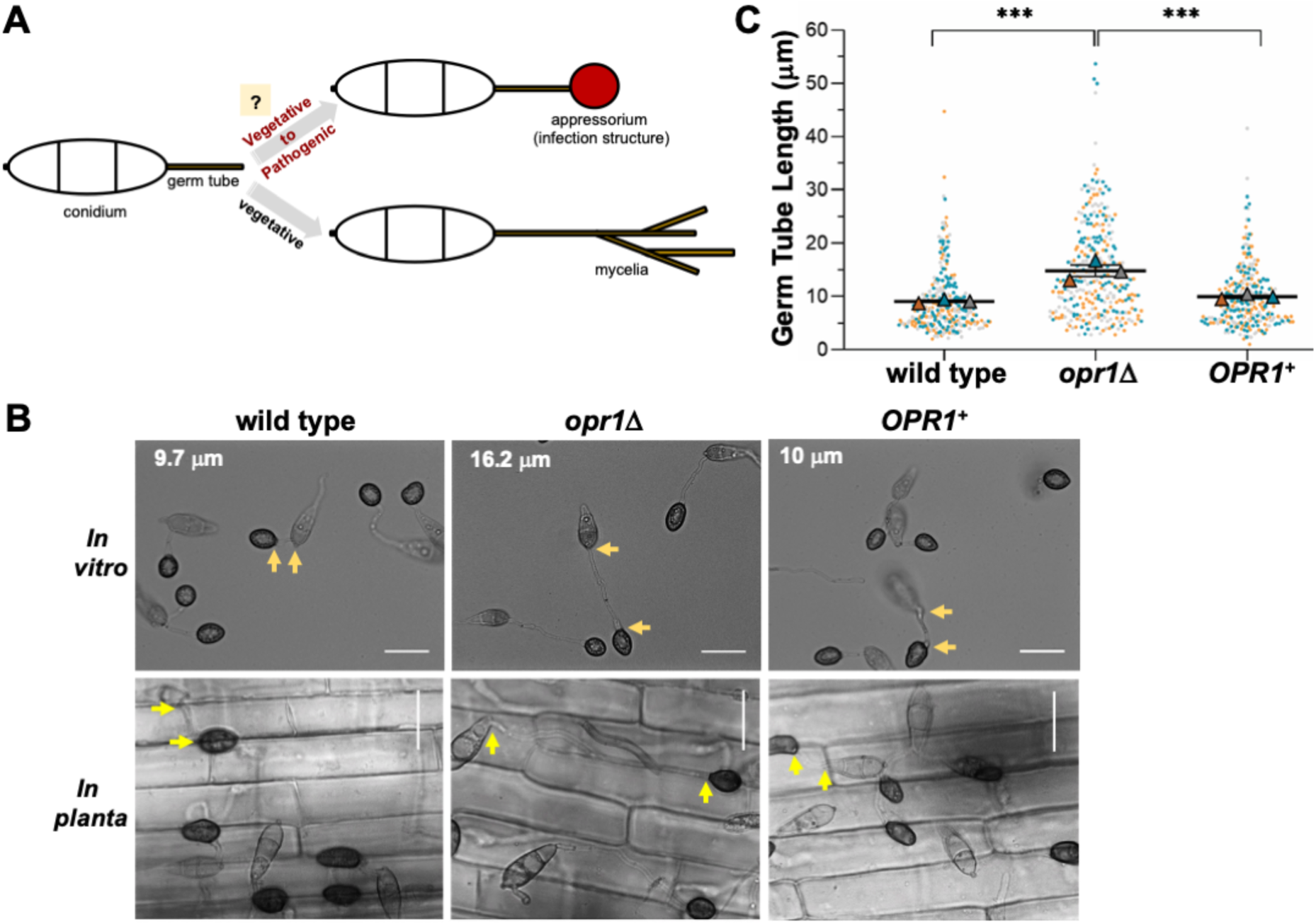
Loss of *OPR1* results in extended germ tube growth prior to appressorium initiation in *M. oryzae*. (**A**) Schematic representation of the critical steps involved in infection-related morphogenesis in conidial germ tubes of *M. oryzae* in response to inductive host environmental cues. The absence of such host factors causes cessation of this developmental switch and continuation of vegetative growth as mycelia without the formation of appressoria. (**B**) Bright-field micrographs showing comparative analysis of germ tube growth and appressorium formation in conidia from wild-type, *opr1*Δ, or the complemented *opr1*Δ (*OPR1^+^*) *M. oryzae* strains inoculated on artificial inductive surface (*in vitro*) or rice sheath (*in planta*). Compared to the WT or Complemented strain, germ tubes in *opr1*Δ mutant were significantly longer. Arrows demarcate the germ tube length in each panel. Scale bar equals 10 μm. (**C**) Graphical representation of the quantification of the germ tube length in the indicated *M. oryzae* strains. Each dot represents the individual data derived from three independent biological repeats experiments (n=300 conidia for each sample). The overlapping triangles represent the means of each repeat in line with dots of the same colour. Two-tailed *t*-test was applied for the comparisons, ***, P<0.001. Fresh conidia were inoculated on hydrophobic cover glass or rice sheath, and measurements performed at 24 hpi.

Here, we demonstrate that the phytopathogenic fungus *M. oryzae* synthesizes jasmonic acid via a fungal OPDA reductase, MoOpr1. Interestingly, loss of *OPR1 significantly* reduced the overall JA levels and increased *cis*-OPDA accumulation *in vivo*. Our study revealed that JA synthesis in *M. oryzae* follows the octadecatrienoic pathway akin to plants, with *cis*-OPDA as the essential precursor moiety. We identify an essential function for fungal oxylipins JA and *cis*-OPDA in appressorium development and function in rice blast, revealing a hitherto unknown regulatory signalling circuit that functions together with G protein cascade in as a chemico-morphogenetic switch enabling pathogenic development and disease in the devastating rice blast fungus.

## Results and Discussion

### Fungal jasmonate is essential for proper cessation of germ tube growth during pathogenic differentiation in *M. oryzae*

Based on the JA biosynthesis pathway and sequence similarity to the orthologous gene (AT2G06050) in Arabidopsis, we predicted *OPR1* (*12-oxophytodienoate reductase 1*, MGG_10583; UniProt accession G5EHQ2; 44.2% similarity to AtOpr1; **Fig. S1A, B**) as a candidate locus for intrinsic jasmonate synthesis in the rice-blast fungus *Magnaporthe oryzae*. To investigate the cellular function, an *opr1*Δ strain was generated by replacing the entire *MGG_10583* ORF with bialaphos-resistance marker cassette in the *M. oryzae* wild-type strain; and a genetically complemented strain also created by introducing the full-length *GFP-OPR1* into the *opr1*Δ **(Fig. S2A, B)**. A vast majority of *opr1*Δ conidia produced significantly longer germ tubes compared to the WT or the genetically complemented *opr1*Δ strain **(Fig. 1B)** under *in vitro* (cover glass) and *in planta* (rice sheath) conditions. Detailed quantification and statistical analyses showed that the average germ tube length in WT, *opr1*Δ, and complemented *opr1*Δ strain was 9.8, 16.0, and 10.0 micron, respectively (**Fig. 1C**), at the time of infection structure formation. Such defects or delay in pathogenic differentiation could be completely suppressed by native expression of GFP-Opr1 in the *opr1*Δ mutant **(Fig. 1B, C)**.

In plants, loss of *OPR* function leads to deficiency in JA production because of a failure to catalyze *cis*-OPDA to 3-oxo-2(2’[*Z*]-pentenyl)cyclopentane-1-octanoic acid, and the resultant defects can be suppressed by exogenous JA but not the precursor *cis*-OPDA (Stintzi 2000, Chehab, Kim et al. 2011). Interestingly, chemical complementation with exogenous jasmonic acid was likewise sufficient to restore proper germ tube development prior to initiation of infection structures in the *opr1*Δ mutant (**Fig. 2A, B**). Likewise, additional JA further shortened the germ tubes in the wild-type strain. Conversely, treatment of WT *M. oryzae* with phenidone, an inhibitor of JA biosynthesis, resulted in significantly elongated germ tubes reminiscent of the *opr1*Δ phenotypic defect of delayed appressorium formation (**Fig. 2C, D**). Taken together, we conclude that fungus-derived intrinsic jasmonic acid is essential for proper and timely cessation of vegetative growth prior to initiation of appressorium formation; and further infer from the chemical complementation and inhibitor analyses that *OPR1* is involved in JA biosynthesis in the blast pathogen.

**Figure 2.**
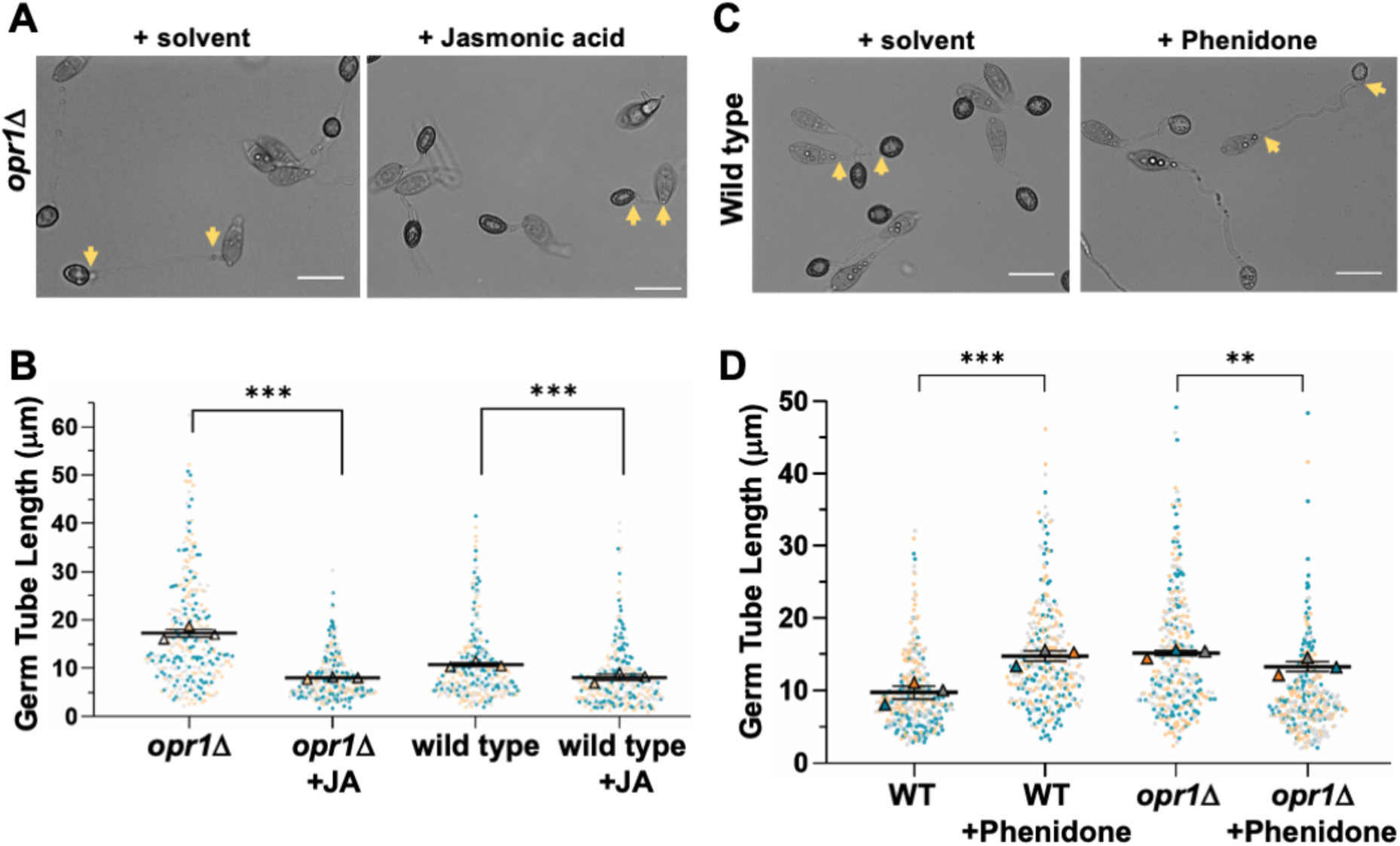
Exogenous JA suppresses germ tube defects in the *opr1*Δ mutant. (**A**) Conidia from the *opr1*Δ strain were inoculated on inductive surface in the presence (0.2 mM) or absence (mock/solvent control) of exogenous jasmonic acid, and germ tube and appressorial development assessed and quantified at 24 hpi. Germ tube length indicated by 2 arrows in each panel. Scale bar = 10 µm. (**B**) Exogenous JA significantly represses germ tube growth in *opr1*Δ and WT strain. Data represents mean±SE from 3 independent repeats of the experiment each involving at least 100 conidia. Each dot represents the value in the data from three biologically independent repeats (n=300). Two-tailed *t*-test was performed to test the differences among the samples, ***, P<0.001. (**C**) and (**D**) Chemical inhibition of JA synthesis (phenidone) results in elongated germ tubes and recapitulates the *opr1*Δ phenotype in the wild-type *M. oryzae* strain. Conidia from the WT strain were inoculated on inductive surface in the presence or absence of the JA-inhibitor Phenidone, and germ tube and appressorial development quantified at 24 hpi. Germ tube length delineated by 2 arrows in each panel. Scale bar = 10 µm. Quantification performed as detailed for panel **B** above.

### Opr1 is required for JA biosynthesis in *M. oryzae*

Based on the chemical complementation experiment, we hypothesized that JA synthesis and accumulation are significantly reduced/altered in the *opr1*Δ strain. To precisely investigate and quantify JA biosynthesis in *M. oryzae*, LC-MS was applied to analyse the overall levels of JA and its major precursor *cis*-OPDA. Targeted metabolite profiling demonstrated that about 34.85 pg/ml fungal JA (m/z 211.10) was produced by the wild-type *M. oryzae* strain. Loss of Opr1 nearly halved the overall accumulation in the *opr1*Δ mutant **(Fig. 3A)**. In contrast, the substrate of Opr1, *cis*-OPDA (m/z 293.00), showed a markedly increased accumulation in the *opr1*Δ mutant, which was estimated to be about five-fold higher than the WT levels **(Fig. 3B)**. Based on such an abnormal accumulation of *cis*-OPDA and a significant reduction in overall JA content in the *opr1*Δ mutant, we construe that Opr1 utilizes OPDA as a substrate and plays an important role in jasmonate/oxylipin biosynthesis in the rice blast fungus.

**Figure 3.**
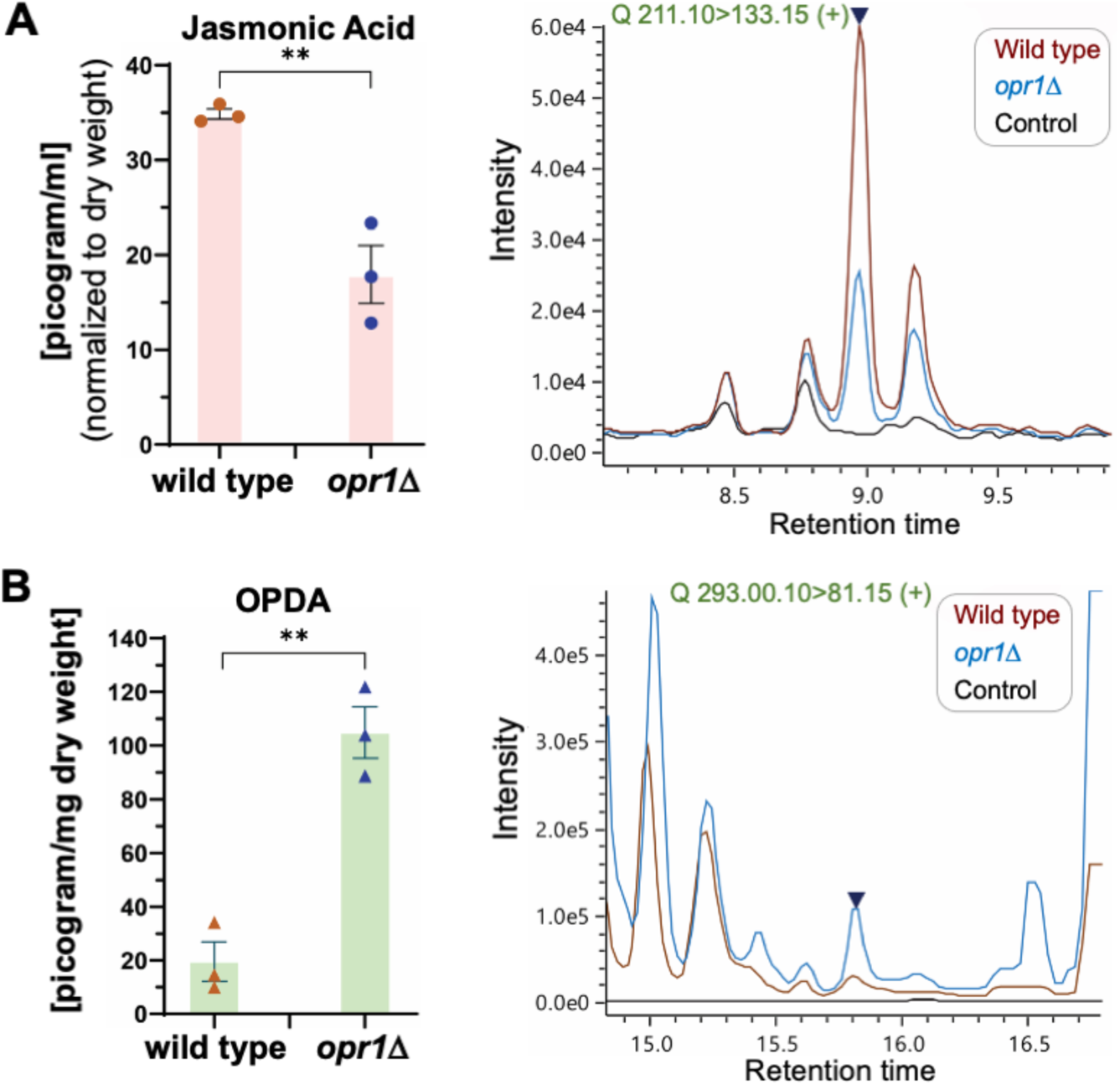
Opr1 is necessary for jasmonate biosynthesis using *cis*-OPDA as a substrate in *M. oryzae*. (**A**) JA production is halved in the oxylipin signature of *opr1*Δ compared to the wild type. Selective ion monitoring for the ion m/z = 211.10 was utilised for detection of fungal JA in wild-type (WT) and *opr1*Δ. Data are the mean ±SE of three repeats of the experiment. (**B**) A significantly higher accumulation of the substrate of Opr1, *cis*-OPDA, occurs in the *opr1*Δ; while only trace levels of the *cis*-OPDA are evident in WT extracts. (**, P<0.01; two-sided *t*-text). Dark blue peak in each spectrum represents the chemical standard used for JA or *cis*-OPDA, respectively. Please refer to the Methods section for more details.

### Subcellular localization of GFP-Opr1 in *M. oryzae*

A GFP-Opr1 expressing strain was generated by in-frame fusion of GFP at the N terminus of the *OPR1* locus to analyze the subcellular localization during asexual and pathogenic development in *M. oryzae* **(Fig. S3)**. The introduction of *GFP-OPR1* restored normal germ tube growth in *opr1*Δ **(Fig. 1B, C)**, thus indicating that GFP-Opr1 is fully functional *in vivo*. GFP-Opr1 signal was found to be distributed predominantly in the cytosol in mycelia and aerial hyphae **(Fig. 4A, B)**, primarily suggesting that GFP-Opr1 is a cytosolic protein. Interestingly, the cytosolic GFP-Opr1 congregated as punctate/vesicular structures in the terminal cell of the conidium, which subsequently forms the germ tube **(Fig. 4C)**. Subsequently, these punctae or vesicles migrated and distributed along the growing germ tube. Upon appressorium formation (8 hpi), the punctate GFP-Opr1 signal diminished; while the ubiquitous cytosolic distribution became more prominent, similar to that observed in mycelia or aerial hyphae. At 24 hpi, the GFP-Opr1 signal was undetectable in the conidium but assembled as punctate structures in the nascent and mature appressorium **(Fig. 4C)**. *In planta*, the mature appressoria form penetrate pegs to initiate the invasive growth. We found that the subcellular localization of GFP-Opr1 at this stage was cytosolic similar to that observed in vegetative hyphae **(Fig. 4D)**. We infer that the punctate or vesicular pool of GFP-Opr1, evident during the early stages of appressorium formation, is indicative of the pivotal role ofOpr1/JA in the precise regulation of germ tube growth concomitant with pathogenic differentiation in rice blast.

**Figure 4.**
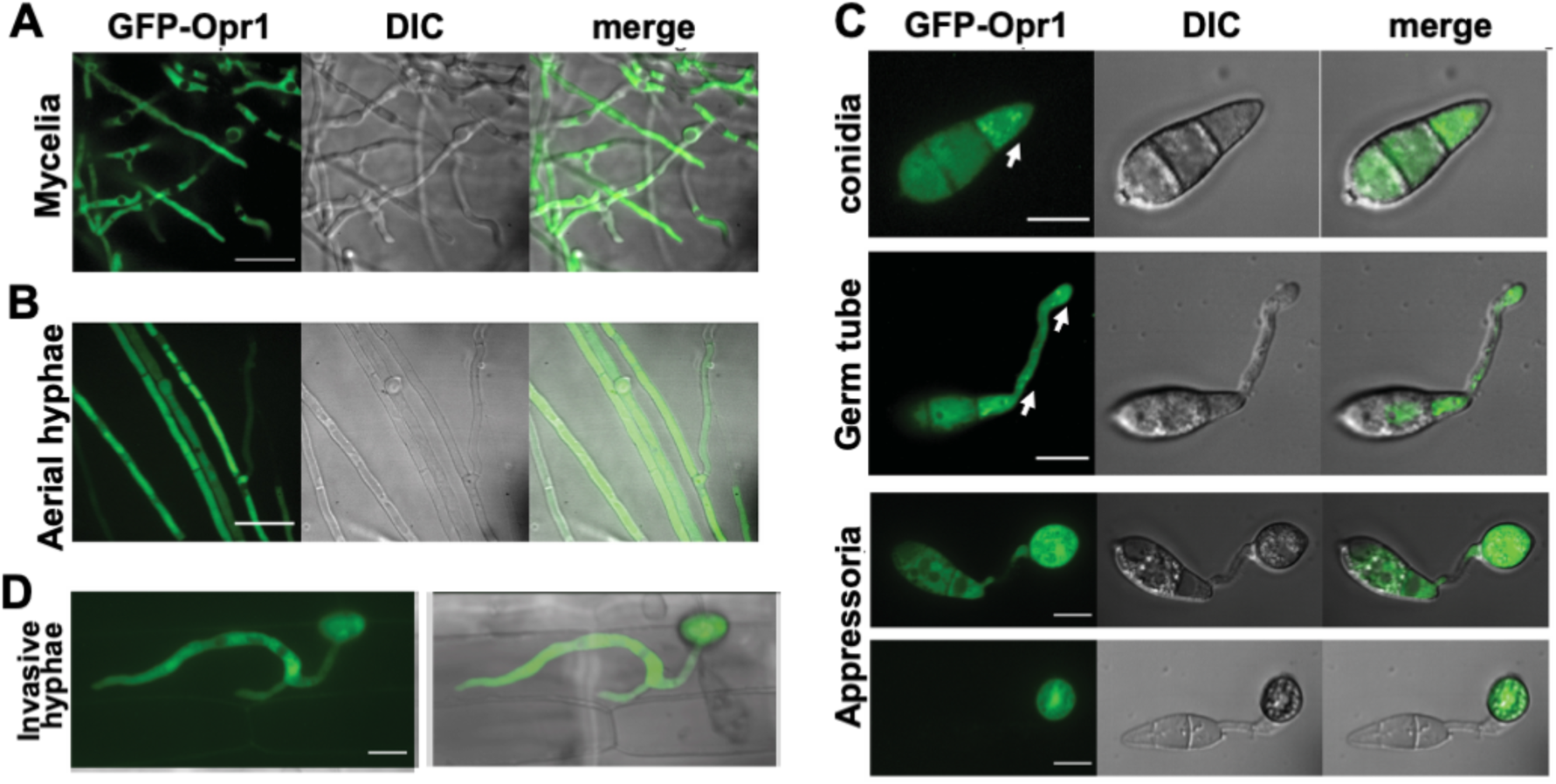
Subcellular localization of the GFP-Opr1 at different stages of vegetative, asexual, and pathogenic development in *M. oryzae*. (**A**) and (**B**) GFP-Opr1 localizes to the cytosol during vegetative growth. Vegetative mycelia and aerial hyphae from the GFP-Opr1 strain were imaged at 3 dpi and 7 dpi, respectively. The size bar equals 10 μm. (**C**) Subcellular localization of GFP-Opr1 in conidia, developing germ tubes, and the incipient appressoria. Arrows indicate the punctate/vesicular localization of GFP-Opr1. Scale bar = 5 μm. (**D**) Subcellular localization of GFP-Opr1 during invasive *in planta* growth of *M. oryzae* in rice. The confocal images were captured at 26-28 hpi, which represents the early invasive growth phase in the host. Scale bar = 5 μm. Images shown are maximum intensity Z-projections of eight confocal stacks, each measuring 0.3 μm.

### Opr1/JA signaling pathway is essential for proper initiation of appressorium formation in *M. oryzae*

Based on the subcellular localization pattern of GFP-Opr1 and the aforementioned characterization of *opr1*Δ, we next asked whether fungal jasmonate plays a role in appressorium formation *per se* in addition to its function in precise cessation of germ tube development. At 4 hpi, 58.2±6.1% WT conidia initiated appressorium formation, whereas only 34.7±1.4% conidia in *opr1*Δ showed the typical hooking stage. Thus, at this early time point, >60% of conidia in the *opr1*Δ mutant failed to initiate infection structures (**Fig. 5A, B**). Exogenous addition of JA caused a significant increase in the ability of such *opr1*Δ conidia to initiate appressorium formation and advanced the time for such induction of pathogenic differentiation to that observed in wild type *M. oryzae* (**Fig. 5A-C**).

**Figure 5.**
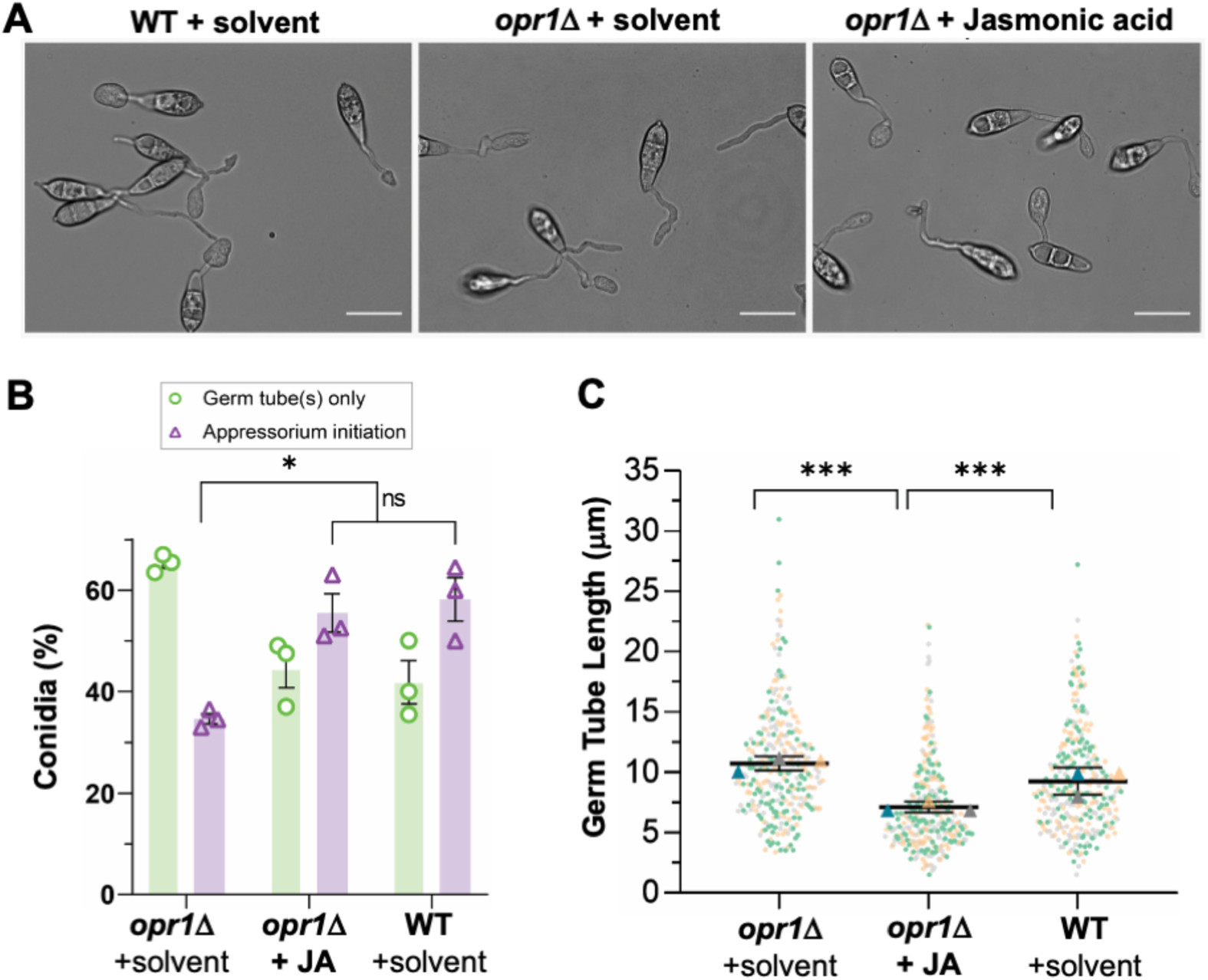
Loss of Opr1 significantly delays appressorium initiation in *M. oryzae*. (**A**) Exogenous JA corrects the delayed appressorium initiation defect in *opr1*Δ. Bright-field micrographs of the developing appressoria in *opr1*Δ and WT strain. Compared to the WT control, overall hooked structures (indicating the initiation of appressoria formation) in *opr1*Δ germ tubes were significantly reduced; but are restored upon treatment with exogenous JA. Conidia were inoculated and grown on the inductive cover glass surface for 4 h. Scale bar 10 μm. (**B**) Quantification of the chemical complementation experiment depicted in A. Data shown represents the mean ±SEM from three independent experiments, and two-tailed unpaired Student’s T-test determined P-values, *, p<0.05. n = 300 conidia per experiment (**C**) Dots and box plot depicting germ tube length in conidia that have initiated appressorium formation in *opr1*Δ and WT strain of *M. oryzae*. Data (n=300) is derived from three independent experiments, with Two-tailed T-test performed to evaluate the significance values, ***, p<0.001.

In order to uncouple such dual function of JA signalling: (i) cessation of vegetative growth (ii) initiation of appressorium formation, we tested the effect of JA on WT *M. oryzae* conidia under a non-inducive condition (soft hydrophilic surface) that does not promote appressorium formation. Under such non-inductive conditions, exogenous JA was found to impede and/or inhibit vegetative growth in germ tubes in a dose-dependent manner (**Fig. 6A**). We conclude that Jasmonic acid signalling is an essential modulator of the vegetative-to-pathogenic switch in the rice blast fungus, and infer that such JA function requires a dual action in first arresting germ development and then promoting appressorium initiation in a precise and timely manner.

**Figure 6.**
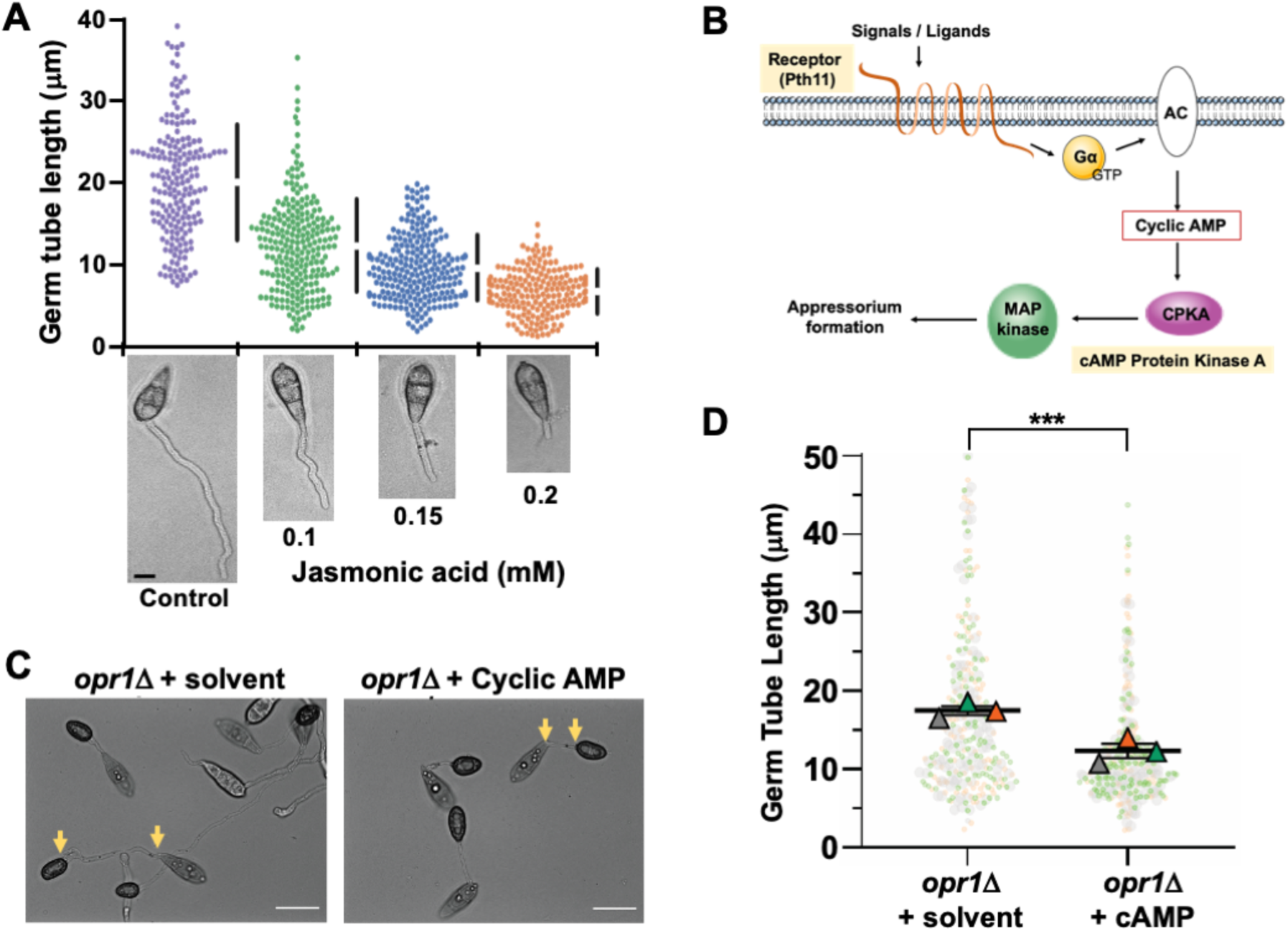
JA regulates germ tube development and appressorium formation in concert with the cyclic AMP signaling in Magnaporthe. (**A**) Exogenous JA shortens the germ tube length/growth on a non-inductive surface. Conidia from the WT strain were inoculated on a non-inductive hydrophilic surface (Gelbond) in the presence of the indicated amounts of JA and germ tube growth assessed at 8 hpi. Data represents mean±SE from 3 independent repeats of the experiment each involving at least 300 conidia. Micrographs of the WT germ tubes formed on non-inductive surface upon treatment with the indicated JA concentration. Scale bar equals 5 μm. (**B**) Schematic depiction of the heterotrimeric G proteins / cyclic AMP-PKA signalling cascade essential for appressorium formation in the rice blast fungus. Exogenous stimuli activate the membrane-localized GPCR Pth11 to trigger the downstream G proteins. Mac1, an adenylate cyclase, catalyses the ATP to cAMP conversion. Induced cAMP during pathogenic differentiation activates the CPKA kinase triggering the downstream signaling, including the Pmk1-MAPK cascade, thus leading to appressorium formation. (**C**) Exogenous cAMP suppresses the dual defects of elongated germ tubes and delayed appressorium formation in the *opr1*Δ mutant. Germ tube development and appressorium formation was assessed on inductive surface in *opr1*Δ conidia in the presence or absence of cyclic AMP. Images were captured at 24 hpi. Scale bar = 10 µm. (**D**) Graphical depiction of the quantitative analysis of the germ tube length in conidia capable of appressorium formation in *opr1*Δ. Dots represent length measurement of each germ tube in three independent replicates of the experiment with n = 300 conidia in each instance. Triangles depict mean±SE value of the three repeats. ***, P<0.001.

### Functional dependency between JA and Cyclic AMP in appressorium initiation in *M. oryzae*

The abnormally elongated germ tubes are a hallmark of mutants defective in cyclic AMP signaling, leading to a delay in or loss of appressorium formation in *M. oryzae* (Xu, Urban et al. 1997, Bindslev, Kershaw et al. 2001, Dąbrowska, Freitak et al. 2009, Kou et al 2017). Therefore, we reasoned that JA/Opr1 function likely cooperates/intersects with the cAMP cascade during appressorium initiation in *M. oryzae*. Since loss of Opr1 delays or disrupts appressorium initiation, we hypothesized that a crosstalk might occur between JA or related oxylipin(s) and the canonical cAMP/MAP kinase signalling, which plays a pivotal role in appressorium development in *M. oryzae* **(Fig. 6B)**. Exogenous cAMP was added to the *opr1*Δ conidia and inoculated on inductive surface to assess the effect on germ tubes and appressorial development. Quantitative analyses at 24 hpi revealed that cAMP shortened the germ tube length from 17.3 μm to 14.0 μm (***P=1.3E-10, two-tailed *t*-test) **(Fig. 6C, D)** in the *opr1*Δ mutant. This led us to infer that a potential functional crosstalk occurs between JA and cAMP pathways during the process of appressorium development in rice blast.

### Functional crosstalk between Jasmonate and cAMP/MAPK signaling during pathogenic development

As shown in Figure 6B, Pth11 is a *bona fide* G-protein coupled receptor (GPCR) that anchors the G-protein/cAMP signalling in Magnaporthe (DeZwaan, Carroll et al. 1999, Ramanujam, Calvert et al. 2013, Kou, Tan et al. 2017). Loss of *PTH11* results in a complete loss of cAMP signalling, leading to highly extended germ tubes that fail to form appressoria. Here, we decided to test whether exogenous JA or the related oxylipins could suppress such *pth11*Δ defects of such elongated germ tubes and lack of appressorium formation. Remarkably, JA or *cis*-OPDA (0.2 mM and 10 μM, respectively) could significantly shorten germ tube growth and advanced the induction of appressorium formation in *pth11*Δ in a significant manner (1.9±1.2% (untreated) to 60.6±8.2% and 22.5±2.9%, respectively) (**Fig. 7A, B**). Furthermore, Methyl JA, a potent inducer of the JA signalling pathway in plants, also restored appressorium formation in *pth11*Δ to 26.7±1.7% (**Fig. 7A, B**) similar to OPDA. In contrast, α-Linolenic acid (α-LeA), the substrate of Lox1 in JA biosynthesis, failed to trigger appressorium formation in *pth11*Δ (**Fig. 7A, B**). To sum up, JA and its derivative (MeJA) and precursor (*cis*-OPDA) but not α-LeA are capable of restoring proper germ tube development and inducing timely appressorium formation in *pth11*Δ, showing that these JA related oxylipins have a similar function downstream of Pth11 in the cAMP pathway to regulate appressorium formation in the blast fungus *M. oryzae*, and JA is the most effective oxylipin (**Fig. 7A**) therein.

**Figure 7.**
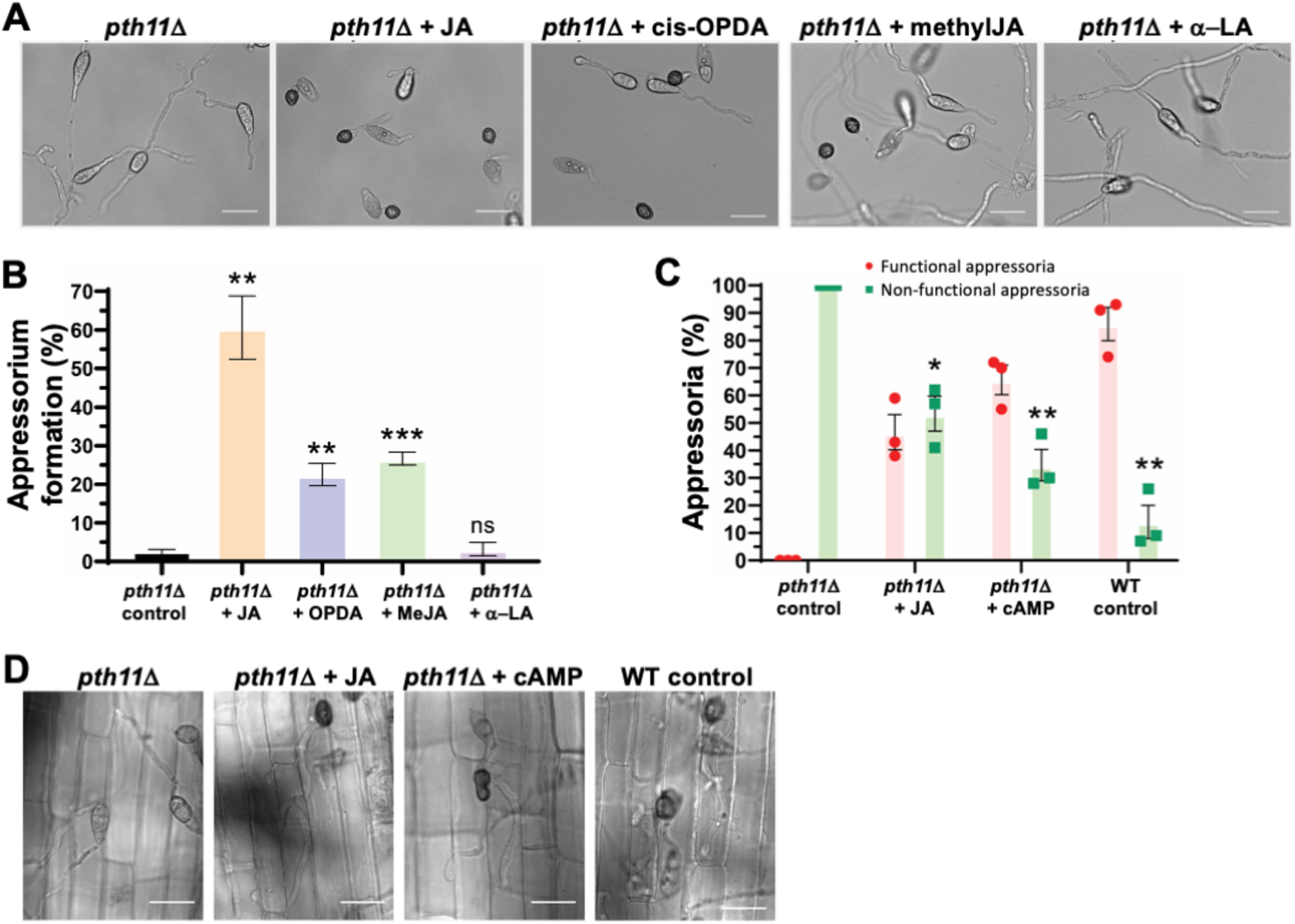
Jasmonate, cis-OPDA and MeJA, but not α-LA, restore appressorium formation and pathogenicity in the *pth11*Δ mutant. (**A**) Exogenous jasmonic acid restores appressorium formation in the *pth11*Δ mutant. Conidia from *pth11*Δ strain were inoculated on the inductive surface for 24 h in the absence or presence of the indicated oxylipin metabolite or *α*-linolenic acid. Treatment with JA, cis-OPDA or MeJA, but not *α*-LA significantly restored appressorium formation in *pth11*Δ. (**B**) Bar graph depicting the quantification of appressorium formation in the *pth11*Δ conidia upon treatment with the indicated oxylipin or precursor fatty acid. Solvent treated *pth11*Δ served as a negative control. (**C**) Oxylipins suppress *pth11*Δ defects in pathogenesis. Bar graph shows that the resultant appressoria in the oxylipin-treated *pth11*Δ mutant are functional, as they were able penetrate the host cells and generate the invasive hyphae at 28 hpi. Treatment with cAMP in parallel or with WT conidia served as positive controls. (**D**) Micrographs showing pathogenic development at 28 hpi in the indicated *M. oryzae* strains. The untreated *pth11*Δ control, which fails to form appressoria or invasive hyphae on rice sheath, served as a negative control. Scale bars represent 10 micron. Data represent mean±S.E from three biological replicates of the experiment using 300 conidia per sample in each instance.

Next, we checked the host penetration ability of JA-triggered appressoria in *pth11*Δ on rice sheath. About 47±11% such appressoria penetrated the host cells and formed invasive hyphae at 28 hpi, indicating that exogenous JA-induced appressoria were, indeed, functional (**Fig. 7C, D**). We conclude that a potential crosstalk occurs between JA and cAMP signalling during pathogenic development in *M. oryzae*, and infer that the JA and cAMP function therein could be reciprocally substituted.

CpkA, the cAMP-dependent protein kinase, functions downstream of the Pth11/G protein signalling **(Fig. 6B)**, and the typical phenotypic defects in appressorium initiation and proper germ tube growth in *cpkA*Δ are similar to those observed in *opr1*Δ (**Fig. 8A**). Therefore, we aimed to understand the mechanistic basis of JA and cAMP crosstalk and addressed whether JA or *cis*-OPDA could restore proper appressorium development in *cpkA*Δ too. Under normal conditions, about 58.2±6.1% conidia from the WT initiate appressorium formation with a hooked structure at the tip of germ tubes at 4 hpi (**Fig. 8B**). Whereas only 16.3.2±2.3% of *cpkA*Δ conidia are capable of initiating appressoria **(Fig. 8A, B)** at this early time point. Exogenous JA remarkably suppressed this delay, and the percentage of appressoria initiation increased to 35.8±3.3% in JA-treated *cpkA*Δ conidia (P=0.004, **P<0.01, two-tailed *t*-test) (**Fig. 8A, B)**. However, only 6.5±1.1% conidia initiated appressoria in the *cis*-OPDA-treated *cpkA*Δ (P=0.014, *P<0.05, calculated by two-tailed unpair *t*-test). Such OPDA-treated germ tubes were short, but most of them did not possess a hooked structure at the tip at this early stage **(Fig. 8A, B)**.

**Figure 8.**
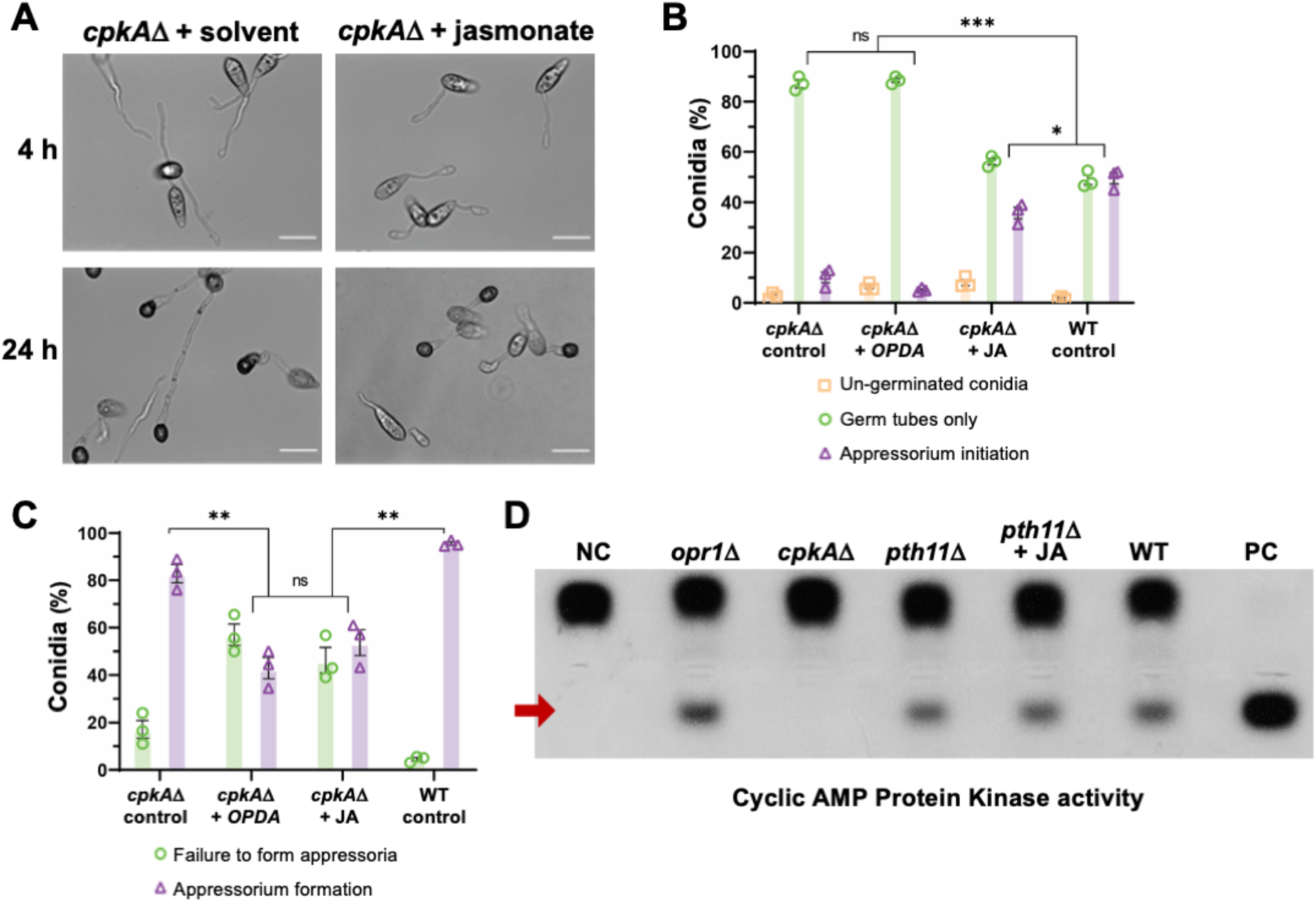
JA signaling functionally interconnects with the cAMP cascade at the CPKA node. (**A**) Exogenous JA, but not *cis*-OPDA, suppresses only the germ tube growth defect in the *cpkA*Δ mutant. Conidia from the *cpkA*Δ strain were treated with JA or *cis*-OPDA and germ tube development and appressorium initiation and formation analysed at 4 hpi and 24 hpi, respectively. Scale bar equals 10 micron. (**B**) and (**C**) JA or *cis*-OPDA fail to restore proper appressorium development in *cpkA*Δ. Bar graphs representing quantitative data analysis for the experiment described in (**A**). Although the oxylipin-treated germ tubes were shorter in length, the percentage of appressorium formation in such JA or *cis*-OPDA treated *cpkA*Δ conidia was significantly reduced. Data represent mean±S.E from three biological replicates each using 300 conidia as sample size. *, P<0.05; **, P<0.01; ***, P<0.001 (unpaired two-tailed t-test; n=3 experiments). (**D**) Loss of *OPR1* or the supplementation with JA does not affect the intrinsic cAMP-Dependent Protein Kinase A (PKA) activity. Quantification of the intracellular cAMP PKA activity in total extracts from the indicated wild-type and mutant *M. oryzae* strains was performed using the non-radioactive CPKA assay system and the model substrate, Kemptide.

**Figure 9.**
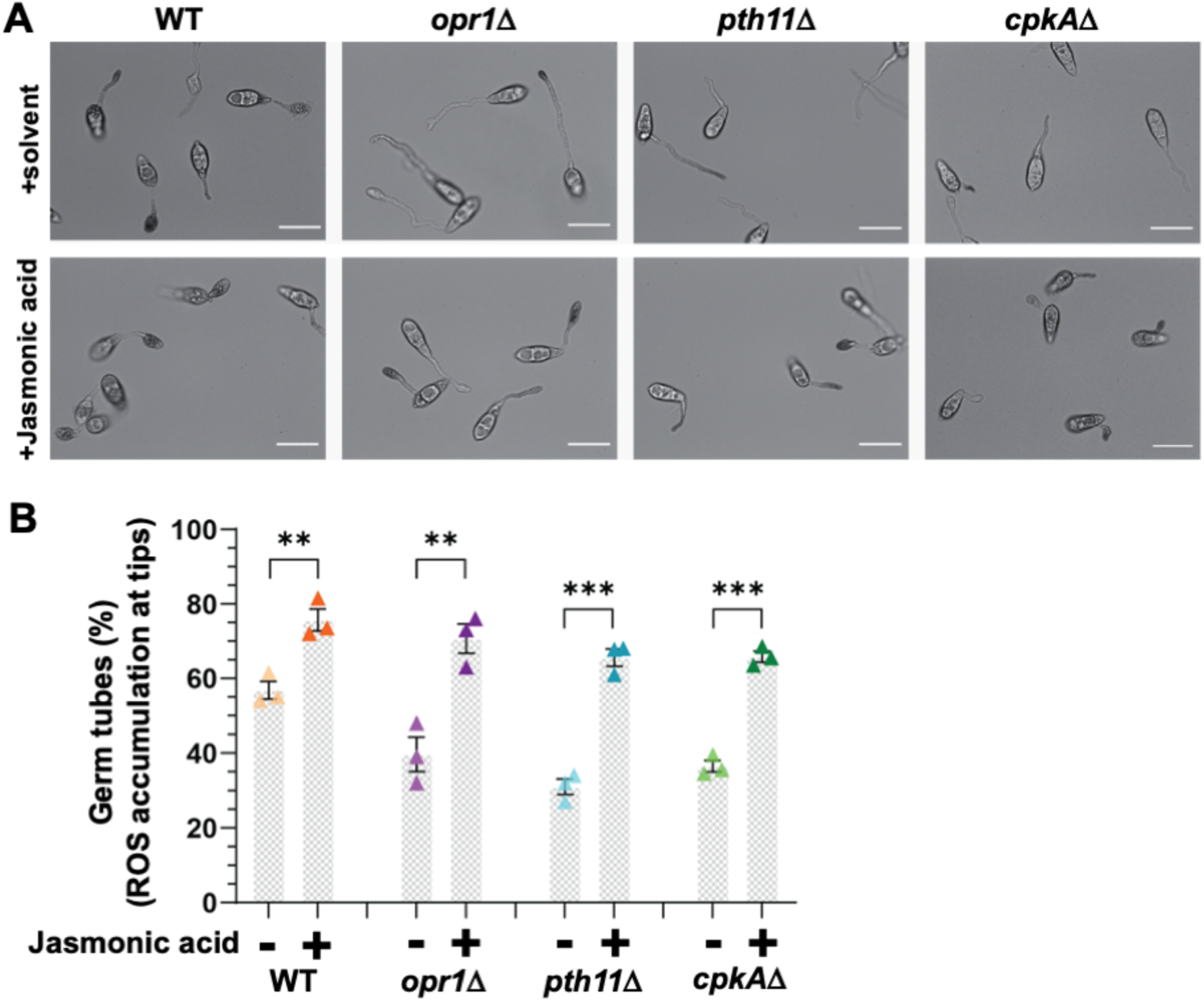
Fungal jasmonate is a key determinant of the cellular redox state during appressorium initiation in *M. oryzae*. (**A**) Loss of Opr1 or the key cAMP regulators (GPCR or CPKA) significantly reduces the overall ROS accumulation; and exogenous JA suppresses such redox defects in the aforementioned mutant strains of rice blast. Conidia from the indicated strains were germinated on inductive surface for 4 h in the presence or absence of 0.2 mM JA and stained with NBT to visualize the ROS at the tips of the germ tubes or hooking structures. (**B**) Bar charts depicting the detailed quantification of ROS accumulations at the germ tube tips of the indicated strains upon treatment with exogenous JA. Data represent mean±S.E from three biological replicates of the experiment each using 300 conidia per sample.

At 24 hpi, treatment with JA or *cis*-OPDA induced the formation of (impaired) appressoria in the *cpkA*Δ conidia. Although the germ tubes were short, the percentage of appressorium formation in JA or *cis*-OPDA treated *cpkA*Δ conidia was significantly reduced from 78.1±9.5% to 49.1±8.4% and 43±6.4%, respectively **(Fig. 8C)**. Neither JA nor cis-OPDA suppressed the morphological defects in cpkAΔ appressoria at 24 hpi. Instead, the percentage of appressoria formed was reduced in the oxylipin treated group **(Fig. 8C)**. A significant number of appressoria in such JA- or OPDA-treated *cpkA*Δ samples were malformed, aberrant or smaller in size compared to the untreated or the WT control groups (data not shown). Furthermore, the overall CPKA enzyme activity was, surprisingly, unperturbed in the *opr1*Δ, and the JA-treated or nascent *pth11*Δ mutant strains (**Fig. 8D**) and was comparable to the WT levels at these time points. Likewise, the levels of active/phosphorylated Pmk1 MAP kinase (MAPK essential for appressorium formation) remained unaltered in the *opr1*Δ mutant (**Fig. S4A**); and exogenous JA failed to restore appressorium formation in the *pmk1*Δ and the *mst11*Δ (MAPKKK) or *mst7*Δ (MAPKK) mutant (**Fig. S4B**)

We conclude that although JA and *cis*-OPDA are able to cause timely cessation of germ tube growth, these 2 oxylipins are incapable of restoring proper appressorium formation in *cpkA*Δ, thus indicating a plausibly complex role for CpkA as a downstream effector in JA signalling in *M. oryzae*. We also infer that the ability of exogenous JA to suppress the *pth11*Δ or *opr1*Δ defects does not involve the utilization and/or induction of CPKA or MAPK activity therein.

### Intrinsic Jasmonic acid modulates redox signaling in M. oryzae

Previously, it has been reported that high levels of ROS accumulate during appressorium development (Egan, Wang et al. 2007, Samalova, Meyer et al. 2014, Kou, Tan et al. 2017). Furthermore, antioxidants such as GSH have been shown to suppress the appressorium formation defect in *pth11*Δ (Kou, Tan et al. 2017). Having ruled out the engagement of MAPK and CPKA activities *per se* as direct downstream effectors of JA function in regulation of germ tube growth, we next addressed whether redox signaling is instrumental in its physiological function(s), since such oxidant responses have recently been shown to be important for cAMP-dependent pathogenic differentiation in *M. oryzae* (Kou et al 2017). Therefore, we assessed and quantified the overall ROS levels in nascent or JA-treated *opr1*Δ and *pth11*Δ by NBT staining. The accumulation of ROS in the germ tube tips was significantly lower in *opr1*Δ*, pth11*Δ, and *cpkA*Δ compared to the wild-type *M. oryzae* (**Fig S4A**). Interestingly, treatment with exogenous JA significantly induced the accumulation of such ROS in the germ tubes termini (**Fig S4A, B**) in the aforementioned 3 mutant backgrounds. Taken together, we infer that a likely mode-of-action for JA-based suppression of germ-tube defects in the *opr1*Δ and *pth11*Δ is to induce and/or buffer the redox homeostasis in these mutant cell types that were found deficient in the overall oxidant capacity. We further construe that JA signalling cooperates with cAMP cascade to determine the cellular redox state during specific stages of infection-related morphogenesis, and plays a key role in modulating the vegetative-to-pathogenic switch in the rice blast fungus.

In conclusion, our study adds to the growing functional importance of oxygenated lipids in fungal biology (Niu, Steffan et al. 2020), and provides new information and insights in implicating fungal jasmonate as a key factor that determines the timely cessation of germ tube growth and precise induction of appressorium formation; and suggests novel cyclic AMP-dependent control points that regulate JA signalling in response to intracellular oxidation in the rice blast pathogen. Future experiments will determine the exact intersection points and mechanistic cooperativity between Jasmonate and Cyclic AMP signaling; and precise biosensor-based real time analysis of fungal JA to reveal novel/critical targets for controlling the devastating blast disease in rice and other important cereal crops.

## Materials and Methods

### Fungal strains, growth conditions and genetic transformation

*Magnaporthe oryzae* strain B157, obtained from the Indian Institute of Rice Research (Hyderabad, India), served as the wild type in this study. For growth and conidiation, wild type and transformants were cultured on prune agar medium (PA; per liter: 40 ml of prune juice, 5 g lactose, 5 g sucrose, 1 g yeast extract, and 20 g agar, pH adjusted to 6.5 with NaOH) or complete medium (CM; per liter: 6 g yeast extract, 6 g casein hydrolysate, and 10 g sucrose and 20 g agar). Assessment of colony characteristics and vegetative growth was carried out by growing the indicated strains in PA or CM medium at 28°C for 3-5 days. For conidiation, fungal strains were cultured in the dark at 28°C for 3 days, followed by continuous light for an extra 5 days at room temperature. Mycelia used for genomic DNA and protein extractions were 2-3 days’ old cultures harvested from liquid CM at 28°C. *Agrobacterium tumefaciens*-mediated transformation (ATMT) was applied to generate fungal transformants. BM (Basal medium; 1.6 g yeast nitrogen base without amino acid and ammonium sulfate, 2 g asparagine, 1 g ammonium sulfate, and 10 g glucose and 20 g agar, pH adjusted to 6.0 with Na_2_HPO_4_) containing 40 mg/ml ammonium glufosinate or chlorimuron ethyl (sulfonylurea, Cluzeau Info Labo, France), and CM with 250 mg/ml hygromycin (A.G. Scientific Inc, USA) was used to select the indicated transformants. *A. tumefaciens* AGL1 was used for T-DNA insertional transformations. Requisite transformants were screened by Southern blot analysis and/or locus-specific PCR, and two confirmed strains selected in each instance for further investigation. *Escherichia coli* strain XL1 was used for bacterial transformations, and maintenance of various plasmids, using specified methods as described (Selvaraj, Shen et al. 2017).

### Nucleic acid manipulations

Homology search of DNA/protein sequences was carried out using NCBI BLAST (Altschul 1997). Multiple protein sequences alignments were performed with ClustalW (Thompson, Higgins et al. 1994), and rendered using Boxshade. Oligonucleotide primers used in this study are listed in Supporting information Table S1. Plasmid DNA was extracted from *E. coli* with Geneaid high-speed plasmids mini kit. Fungal genomic acid DNA was isolated with the MasterPure^TM^ Yeast DNA Purification Kit (Lucigen, Wisconsin, USA) following the manufacturer’s instructions. Nucleotide sequences were analyzed by the ABI Prism Big Dye Terminator Method (PE Applied Biosystems).

### Generation of constructs for gene deletions or epitope tagging

For gene deletion of *OPR1*, DNA fragments (~1 kb) of 5’ UTR and 3’ UTR were PCR amplified, digested, and ligated sequentially to flank the *BAR* resistance cassette in pFGL822-2. GFP-Opr1 construct, the N terminal tagging with the native promoter, was created by fusing GFP fragment with *OPR1* ORF at its C terminal and downstream of its 1kb promoter, sequentially cloned into pFGL1010. This plasmid had a sulfonyl urea resistance and contributed to ectopic single-copy integration by introducing the gene into the ILV2 locus (Yang and Naqvi 2014). Primers used in the plasmid construction were listed in Supporting information table S1. The gene deletion or epitope tagging constructs were introduced into *M. oryzae* B157 via *Agrobacterium*-mediated transformation to replace the target genes specifically.

### Appressorial assays and chemical complementation analyses

For appressorial assays, conidia were collected from a 7-day old PA plate by scraping with an inoculation loop in the presence of sterile water. The conidial suspension was obtained by filtering through two layers of Miracloth (Calbiochem, San Diego, USA), centrifuging and re-suspending in sterile water at a required concentration (106 conidia per ml). Conidial droplets (~20 μl) were placed on the indicated surfaces and incubated under humid conditions at room temperature. Appressoria initiation/formation was observed and quantified at 4h, 24h, and 28h post-inoculation (hpi), denoted in the legends. Appressoria restorations ware carried out by adding Jasmonic acid (JA, Sigma-Aldrich, Darmstadt, Germany), phenidone (PHEN, Sigma-Aldrich, Darmstadt, Germany), 12-oxo Phytodienoic Acid (*cis*-OPDA, Cayman, Ann Arbor, USA), Methyl jasmonate (MeJA, Sigma-Aldrich, Darmstadt, Germany), α-Linoleic acid (α-LA, Sigma-Aldrich, Darmstadt, Germany), Sig8-bromoadenosine 3′,5′-cyclic monophosphate sodium salt (8-Br-cAMP, Sigma-Aldrich, Darmstadt, Germany) at 0 hpi to a final concentration of 0.2 mM, 0.3 mM, 10 μM, 0.5 mM, 0.5 mM, and 10 mM, respectively (Ramanujam, Calvert et al. 2013, Kou, Tan et al. 2017).

### Real time qRT-PCR

Total RNA was isolated from developing appressorial inoculated on hydrophobic glass slides (100 Deckglaser, Thermo Scientific, Germany) at the specified time points with TRIzol LS Reagent (Thermo Fisher, CA, USA) following the manufacturer’s instruction. About 500 μg RNA, each sample was used for the cDNA synthesis carried out by the iScript cDNA synthesis kit (BioRAD, CA, USA). The real-time PCR reactions were performed by the KAPA SYBR FAST qPCR Master Mix Kit (KAPABIOSYSTEMS, Massachusetts, USA), and the requisite primers used in qRT-PCR are presented in Table S1. All qRT-PCR reactions were repeated three times, with three replications for each sample. The cycle threshold (ΔCt) of each gene was normalized against the transcript level of *TUBULIN* (*MGG_00604*).

### Protein isolation and western blot analysis

Total protein was extracted from 2-3 days CM cultured mycelia, which were ground to fine powder with liquid nitrogen, and re-suspended in protein extraction buffer (10 mM Tris-Cl, pH 7.5; 150 mM NaCl; 50 mM NaF; 0.5% NP40; 0.5 mM EDTA; 1 mM PMSF and 1X Protease Inhibitor Cocktail) (Ramanujam, Calvert et al. 2013). About 20 ug total proteins in each sample were fractionated by 10% SDS PAGE gel and transferred to PVDF membranes (Millipore Corporation, USA) and immunoblotted with primary antibody anti-GFP (Invitrogen-Molecular Probes, Thermo Fisher Scientific, USA). For detection, second antibodies IRDye680 conjugated anti-rabbit was used to present the targeted proteins by Odyssey Infrared Imaging System (LI-COR, Lincoln, USA) (Ng, Ng et al. 2020). Coomassie blue staining served as loading control in this study.

### Chemical analysis (Liquid Chromatography-Mass Spectrometry)

Germlings were harvested after growth for 48 hours in liquid Complete Medium, and washed thrice with pre-chilled PBS. The recovered fungal biomass was then frozen immediately at −80 °C and lyophilized and ground to a fine powder using a mortar pestle and liquid nitrogen. This powdered biomass was immediately stored at −80 °C until further use.

Lipids were extracted from 100 mg of ground germlings overnight at 4 °C with 4.5 mL of methanol (MeOH): water (H_2_O) (25:75 by v/v) at 750 rpm (revolution per minute) in a thermomixer (Eppendorf, Hamburg, Germany). Lipids from 100 µl of conditioned media were extracted by vortexing for 2 min with 4.5 mL of methanol (MeOH): water (H_2_O) (25:75 by v/v).

Extracts were centrifuged at 4 °C for 10 min at 4000 rpm in an Eppendorf 5810R centrifuge, and supernatant transferred into new 15 ml Eppendorf tubes, and topped up to a final volume of 6 ml with methanol (MeOH): water (H_2_O) (25:75 by v/v).

Samples were spiked with 50 μl of a deuterated internal standard solution containing PGD2-d4 and PGE2-d4. Analytes were extracted using Strata-X 33 μm Polymeric solid reversed phase (SPE) extraction columns (8E-S100-TGB, Phenomenex). Columns were conditioned with 3 mL of 100% MeOH and then equilibrated with 3 mL of H_2_O. After loading the sample, the columns were washed with H_2_O: MeOH (75:25 by v/v) to remove impurities, and the metabolites were then eluted with 1.5 of 100 % MeOH. The eluant was dried under vacuum and resuspended in 70 μl of ACN/water/formic acid (10/90/0.1, v/v). The extracted samples were then subjected to mass spectrometry analysis.

High-performance liquid chromatography (HPLC) coupled to triple-quad mass analysis was performed by using the Shimadzu LCMS-8060 system. Reversed phase separation was performed on a Phenomenex, Kinetex C8 (2.1 × 100 mm I.D × 150 mm L., 2.6 µm) column and maintained at 40 °C. The mobile phase consisted of (A) water/formic acid (100/0.1, v/v) and (B) ACN. The stepwise gradient conditions were carried out for 30 min as follows: 0 min, 10% of solvent B; 0–5 min, 10–25% of solvent B; 5-10 min, 25–35% of solvent B; 10-20 min, 35-75% of solvent B; 20-20.1 min, 75-98% of solvent B; 20.1-28 min, 98% of solvent B; 28-28.1 min, 98-10% of solvent B; and final 28.1-30 min, 10% of solvent B. The flow rate was 0.4 mL/min, injection volume was 10 µl, and all samples were maintained at 4 °C throughout the analysis.

A mixture of representative native and internal standards was injected and run with the column to optimize the source parameters. The electrospray ionization was conducted in positive mode. Drying gas temperature was set at 270°C with a gas flow of 10 L/min. Sheet gas temperature was set at 250 °C with a gas flow of 10 L/min. The nebulizer gas flow was 230 kPa. The dynamic MRM option was used and performed for all compounds with optimized transitions and collision energies. MRM transitions (precursor and product ions) and collision voltages were as follows: *cis*-OPDA (293 → 81.15; −30 eV), Jasmonic acid (211.1 → 133.15; −13 eV). The determination and integration of all peaks was manually performed using the LabSolutions Insight software. Peaks were smoothed before integration and peak to peak Signal/Noise ratios were determined using the area under the peaks.

### Nitro Blue Tetrazolium staining

Conidial suspension was inoculated on artificial hydrophobic coverslips with or without the indicated chemicals. At 4 hpi, the droplets were removed and replaced with an equal volume of 0.1% Nitro blue tetrazolium (NBT; Sigma-Aldrich, Darmstadt, Germany) solution, staining for 20 mins. The staining was stopped using 100% ethanol, and the samples washed with water three times before the imaging process.

### Live cell microscopy and image analyses

Time-lapse or live-cell fluorescence microscopy was performed using a Zeiss Axiovert 200 M microscope (Plan Apochromat 1006, 1.4NA objective) with an Ultra-View RS-3 spinning disk confocal system (PerkinElmer Inc., United States) which equipped with a CSU21 confocal optical scanner, 12-bit digital cooled Hamamatsu Orca-ER camera (OPELCO, Sterling, VA, USA) and a 491 nm 100 mW and a 561 nm 50 mW laser illumination under the control of MetaMorph Premier Software (Universal Imaging, USA) (Ramanujam, Calvert et al. 2013, Selvaraj, Tham et al. 2017). Briefly, z-stacks comprised of 0.5 µm-spaced sections that were captured. GFP excitation were performed at 491 nm (Em. 525/40 nm).

Bright-field microscopy was carried out by an Olympus IX71 microscope (Olympus, Tokyo, Japan) equipped with a Plan APO 100X/1.45 objective. Images were captured by Photometrics CoolSNAP HQ camera (Tucson, AZ, USA) and processed with MetaVue (Universal Imaging, Downingtown, USA), Adobe Illustrator (Adobe Inc, USA), and ImageJ (LOCI, University of Wisconsin, USA).

### Plant cultivar and pathogenicity assays

Rice cultivar CO39 susceptible to *M. oryzae* strain B157 was utilized for pathogenicity assays. Rice seeds were soaked in water and placed at room temperature for 5 days to germinate. The rice seedlings were grown at 80% humidity, 28 °C in a 16-h light (28 °C)/8-h dark (22°C) cycle, for 2 weeks. For blast infection assays, fresh conidial suspension (1×10^6^ conidia/ml with 0.01% gelatin) was sprayed on the rice seedlings, and inoculated seedlings grown for an extra 7 days in a growth chamber (22 °C, 80-95% humidity, 16 h illumination per day) and the assessment of blast disease symptoms conducted as described (Deng, Qu et al. 2012).

## Acknowledgements

We thank Thomas Dawson and Sanjay Swarup for facilitating the collaborations for lipidomics analyses, and Peter Benke (Singapore Lipidomics Incubator) for help with the initial mass spectrometry analysis. We are grateful to the Fungal Patho-Biology Lab (TLL, Singapore) for helpful discussions and suggestions. This research was supported by grants from the Temasek Life Sciences Laboratory, Singapore, and the National Research Foundation (NRF-CRP16-2015-04), Singapore.

